# Sexual Dimorphism in Systemic Inflammatory Responses to Femur Fracture in Mice Infected with SARS-CoV-2-Like Virus

**DOI:** 10.1101/2023.12.04.567060

**Authors:** Matthew Patrick, Austin Foster, Arun Aneja, Ramkumar T. Annamalai

## Abstract

Patients with femur fractures who are concurrently infected with COVID-19 face a threefold increase in mortality, likely due to a compounded inflammatory response. Furthermore, sex-specific differences in immune responses to COVID-19 have been documented, implicating gender as a potential modulator of disease severity in these comorbid conditions. Understanding the inflammatory interplay underlying this association is critical for the development of effective, targeted therapies to mitigate mortality.

In this study, we investigated the systemic, sex-specific inflammatory response in mice that sustain a fracture while infected with a murine coronavirus (MHV), which belongs to the same genus as SARS-CoV-2. Our findings reveal that the combined inflammatory incidents of MHV infection and fracture disrupt the systemic immune response in both female and male mice, leading to immune dysregulation characterized by altered cell recruitment and disruption of the normal inflammatory cascade. Notably, the study identifies sex-specific differences in immune response, with female subjects exhibiting significantly elevated levels of inflammatory cytokines, including IL-18 and TNFα, while males exhibit a diminished response. These sexually dimorphic differences are also reflected in the systemic immune cell populations, suggesting that the quantity of immune factors released may contribute to the observed discrepancies. Notably, these differences were minimal or moderate in animals that either got an MHV infection or fracture alone.

Our findings indicate that the overproduction of proinflammatory cytokines, such as IFNγ, IL-18, and TNFα—reminiscent of cytokine storm syndrome—drives immune dysregulation, exacerbating outcomes in patients with these comorbidities. The observed sex-specific responses may be influenced by factors such as sex hormones, including estrogen, highlighting the importance of considering gender in therapeutic approaches. These insights provide a foundation for the development of tailored interventions to improve outcomes for COVID-19 patients with musculoskeletal trauma, including fractures.

## 1. INTRODUCTION

The COVID-19 pandemic, caused by SARS-CoV-2, has had an unparalleled global impact, infecting over 650 million individuals and resulting in more than one million deaths in the United States alone^1^. Although the public health emergency officially ended in May 2023, COVID-19 remains a persistent health challenge, transitioning into an endemic phase with long-term consequences still under investigation. Approximately 30–40% of patients report symptoms associated with post-COVID-19 syndrome, or “long COVID,” and the virus continues to exacerbate pre-existing comorbidities, significantly increasing mortality rates^2^.

COVID-19 pathogenesis typically unfolds in two phases: an initial phase driven by viral replication, followed by a secondary phase dominated by the host’s immune response^3^. In severe cases, the immune response can become dysregulated, resembling sepsis-induced cytokine release syndrome (CRS)^4^. This hyperinflammatory state often results in extensive tissue damage and acute respiratory distress syndrome (ARDS), which is a primary cause of mortality in COVID-19 patients^5^. Although immune dysregulation occurs in both sexes, males are more susceptible due to a dampened immune respons^6^. Conversely, females tend to exhibit more robust immune activity, which, when coupled with comorbid conditions, may increase their susceptibility to CRS or ARDS^7^.

Sexual dimorphism in immune responses is a well-documented phenomenon, with males and females exhibiting distinct differences in immune system function^8^. Females typically demonstrate heightened innate and adaptive immune responses compared to males, which may confer greater protection against some pathogens but also increase their risk of developing hyperinflammatory conditions, such as CRS, in response to immune challenges. These differences are influenced by a complex interplay of genetic, hormonal, and environmental factors. Estrogen, for example, enhances immune cell activation and cytokine production^9^, while testosterone generally exerts immunosuppressive effects^10^. This hormonal influence is particularly relevant in the context of COVID-19^11^, where sex hormones may modulate the severity of both the infection and its associated comorbidities.

Sexual dimorphism extends beyond immune function to musculoskeletal tissues, influencing development, repair, and regeneration^12^. Some studies suggest that males exhibit faster fracture healing rates than females, potentially due to lower systemic inflammatory responses^12, 13^. However, this advantage may diminish under conditions of systemic inflammation, such as those induced by COVID-19. Conversely, the heightened immune activity in females could amplify the inflammatory response triggered by fractures, potentially exacerbating complications such as cytokine storms or ARDS. Despite these findings, the underlying mechanisms driving sex-based differences in fracture healing and systemic inflammation remain inadequately understood.

Comorbidities such as diabetes^14^, hypertension^15^, and fractures^16^ exacerbate the severity of COVID-19, compounding the inflammatory burden and complicating recovery. Among these, traumatic musculoskeletal injuries, particularly fractures, pose a unique and poorly understood risk. Clinical evidence indicates that proximal femoral fractures elevate mortality rates in COVID-19 patients from 10.3% to 30.4%^17^, suggesting a synergistic interaction between fracture healing and the immune dysregulation triggered by SARS-CoV-2 infection. A deeper understanding of this intersection, particularly in the context of sex-specific differences, is essential for developing targeted therapeutic strategies to mitigate the elevated mortality associated with these conditions.

Animal models provide critical insights into the complex interactions between COVID-19 and fracture-induced inflammation. In this study, we employed murine hepatitis virus (MHV), a coronavirus of the same genus as SARS-CoV-2, which is widely used to model the inflammatory cytokine responses seen in human SARS-CoV-2 infection^18, 19^. The MHV model closely recapitulates the inflammatory dynamics observed in COVID-19 patients, allowing for controlled investigation of the immune response to fractures in the context of viral infection, while minimizing the biosafety risks associated with SARS-CoV-2.

Here, we explore the systemic immune response to concurrent fracture and viral infection, emphasizing sex-specific differences. By examining the interplay between fracture healing and viral-induced inflammation in male and female mice, this study aims to unravel the sexually dimorphic immune mechanisms that contribute to the heightened mortality observed in these comorbid conditions. Our findings provide critical insights into the pathophysiology of these interactions and lay the groundwork for tailored therapeutic strategies to improve outcomes in patients facing the dual challenges of COVID-19 and musculoskeletal trauma.

## 2. MATERIALS AND METHODS

### 2.1. Animals and studies

All animal procedures were performed in compliance with the protocol approved by IACUC at the University of Kentucky. 12-week-old, wild-type C5BL/6 mice were randomly assigned to either control (n = 10, 5 male and 5 female), fracture (FX) (n = 10, 5 male and 5 female), MHV (n = 10, 5 male and 5 female), or MHV + fracture (MHV+FX) (n = 10, 5 male and 5 female) groups. MHV mice were infected intranasally inside an ABSL2 facility with 10^4^ PFU murine hepatitis virus (MHV-VR764, ATCC), as shown previously^18, 19^. 48 hours after the intranasal inoculation, the mice in fracture groups were subjected to transverse diaphyseal fracture using a well-established blunt-trauma method as described^20^. Briefly, mice were anesthetized using ketamine (1.5 mg/kg of body mass) and xylazine (1.5 mg/kg of body mass), a bland ophthalmic ointment was applied, and the surgical site was shaved and wiped with 70% ethanol, followed by 10% povidone-iodine solution. A small incision was made at the patella, and it was laterally dislocated to expose the knee joint. Using a 0.5 mm centering bit, a hole was made in the intercondylar notch, and an intramedullary guide tube (0.2 mm NiTi tube) was inserted into the femur. With the tube in place, the right leg was positioned under the guillotine of a murine fracture device (RIsystems). The blunt striker was then released to produce impact kinetic energy (*E*_*k*_) of ∼40 *mJ*, which is shown to create a transverse fracture in adult C57BL/6 mice^21^. The guide tube was then replaced with a 16 mm titanium intramedullary nail, and the fracture stabilized. The skin was sutured with a 6-0 vicryl suture (Ethicon). Then, 1 mL of saline was administered intraperitoneally for hydration, followed by subcutaneous administration of buprenorphine (1.5 mg/kg body mass) for analgesia. The animals were then placed into a recovery cage and monitored per the established post-operative animal care protocols. Animals were monitored daily for 7 days and weekly thereafter to ensure post-operative recovery.

Blood was collected longitudinally in dipotassium (K2) EDTA microvette tubes (Sarstedt) from the right submental vein at 5 days prior to fracture induction for a baseline and at days 2 and 7 post-induction. Extracted blood was centrifuged at 900 g for 5 minutes to separate plasma, which was then collected and stored at -80°C until analysis. The remaining plasma-free blood samples were immediately stained and analyzed through flow cytometry for different immune cell populations (See section 2.3 for details).

### 2.2. MicroCT imaging and morphometric analysis

The fractured femurs retrieved at the end of our study were scanned using a high-resolution small animal micro-computed tomography (microCT) system (SkyScan1276 microCT, Bruker, Billerica, MA) with an Al 1mm filter with the voltage set to 70 kV and image pixel size 20.32 μm. Implants were carefully removed, and femurs with surrounding musculature were placed in a new dry tube before scanning. Images were then reconstructed (NRecon) with smoothing set to 3, ring artifact correction set to 4, and beam hardening correction set to 30%.

### 2.3. Flow cytometric Analysis

After plasma separation, red blood cells (RBC) were lysed using 1X RBC lysis buffer (eBioscience). Lysis buffer was washed away with Dulbecco’s phosphate-buffered saline (PBS, Gibco), and the cells were centrifuged at 400 G centrifugation for 5 minutes. The cell pellet was fixed with 4% buffered formalin for 15 minutes, washed with PBS, and centrifuged again. The pellet was then suspended in a blocking solution consisting of FACS buffer (1X PBS with 5% fetal bovine serum (Gibco) and 0.1% sodium azide) and anti-mouse CD16/32 (eBioscience) for 10 minutes at 4°C to block Fc receptors. Cells were stained for 45 minutes in the dark at 4°C with various fluorescently labeled anti-mouse antibodies to identify specific immune cell populations. Including lymphoid (CD45^+^/CD11b-), myeloid (CD45^+^/CD11b^+^), M1 and M2 monocytes (Ly6G-/Ly6C^hi/low^), macrophages (Ly6G-/Ly6C^+^/F4/80^+^), M2 macrophages (Ly6G-/Ly6C^+^/F4/80^+^/CD206^+^), dendritic cells (CD11c^+^), T cells (CD3^+^), T helper cells (CD3^+^/CD4^+^), cytotoxic T cells (CD3^+^/CD8^+^), T regulatory cells (CD3^+^/CD4^+^/CD25^+^), and B cells (CD19^+^). All staining solutions were prepared in FACS buffer. Details of antibody conjugates, clones, dilution factors, and suppliers are provided in Supplementary Table 1. After staining, samples were washed and resuspended in FACS buffer, then analyzed immediately using a 4-channel flow cytometer (BD FACSymphony A3). Data were processed, and cell populations were quantified using FlowJo software. Fluorescence-minus-one (FMO) controls for each antibody were employed to establish negative gates, ensuring minimal noise (<1%).

### 2.4. Multiplex Assay

A 48-plex Mouse ProcartaPlex™ Panel (Thermo Fisher, EPX480-20834-901) was used to analyze the plasma isolates from the blood samples. Capture beads were added to each well of the plate, which was subsequently washed using a Hand-Held Magnetic Plate Washer (EPX-55555-000). Following the wash, samples, standards, and low and high controls were added to the wells in duplicate. The plate was then sealed with the provided plate sealer and lid, mixed at a speed of 600 rpm for 2 hours at room temperature, and was washed thrice using the provided wash buffer. A detection antibody mix was added, and the plate was resealed and mixed for 30 minutes at room temperature. The plate was then washed thrice, and Streptavidin-PE was added, resealed, and agitated for 30 minutes at ambient temperature and washed thrice. After adding the reading buffer and incubating for 5 minutes, the plate was processed on an xMAP INTELLIFLEX system with a 50 μL acquisition volume, a DD Gate ranging from 4000 to 13000, and a low PMT reporter gain setting.

### 2.5. Statistics Methods and Analyses

All measurements were performed with at least 4 replicates. Data are plotted as means with error bars representing the standard deviation. A two-way ANOVA was used for statistical analysis of multiple groups, followed by a Holm-Sidak test post-hoc test for multiple comparisons. All statistics analyses were done using GraphPad Prism (version 10.0.3). Differences with *p* < 0.05 were considered statistically significant. Principal component analyses (PCA) were performed in R (version 4.4.0) using PCAtools (version 2.5.15), with data imputed for missing values at day 3 using impute (version 1.78.0). The Multiplex Assay data was excluded from the PCA analysis due to a number of undetected values which may introduce significant variability, potentially skewing the results.

## 3. RESULTS

### 3.1 Fractures Increase Mortality in MHV-Infected Mice

In this study, adult male and female C57BL/6 mice were randomly assigned to four groups: Control (no inoculation or fracture), FX (no inoculation with fracture), MHV (inoculation without fracture), and MHV+FX (both inoculation and fracture). The respective procedures were performed, and all animals were closely monitored for signs of distress and adverse conditions. A schematic of the study design and timeline of procedures is depicted in (**Fig.1A**). A representative microCT image of the mid-diaphyseal fracture induced through blunt trauma is shown in **Fig.1B**.

**Figure 1.**
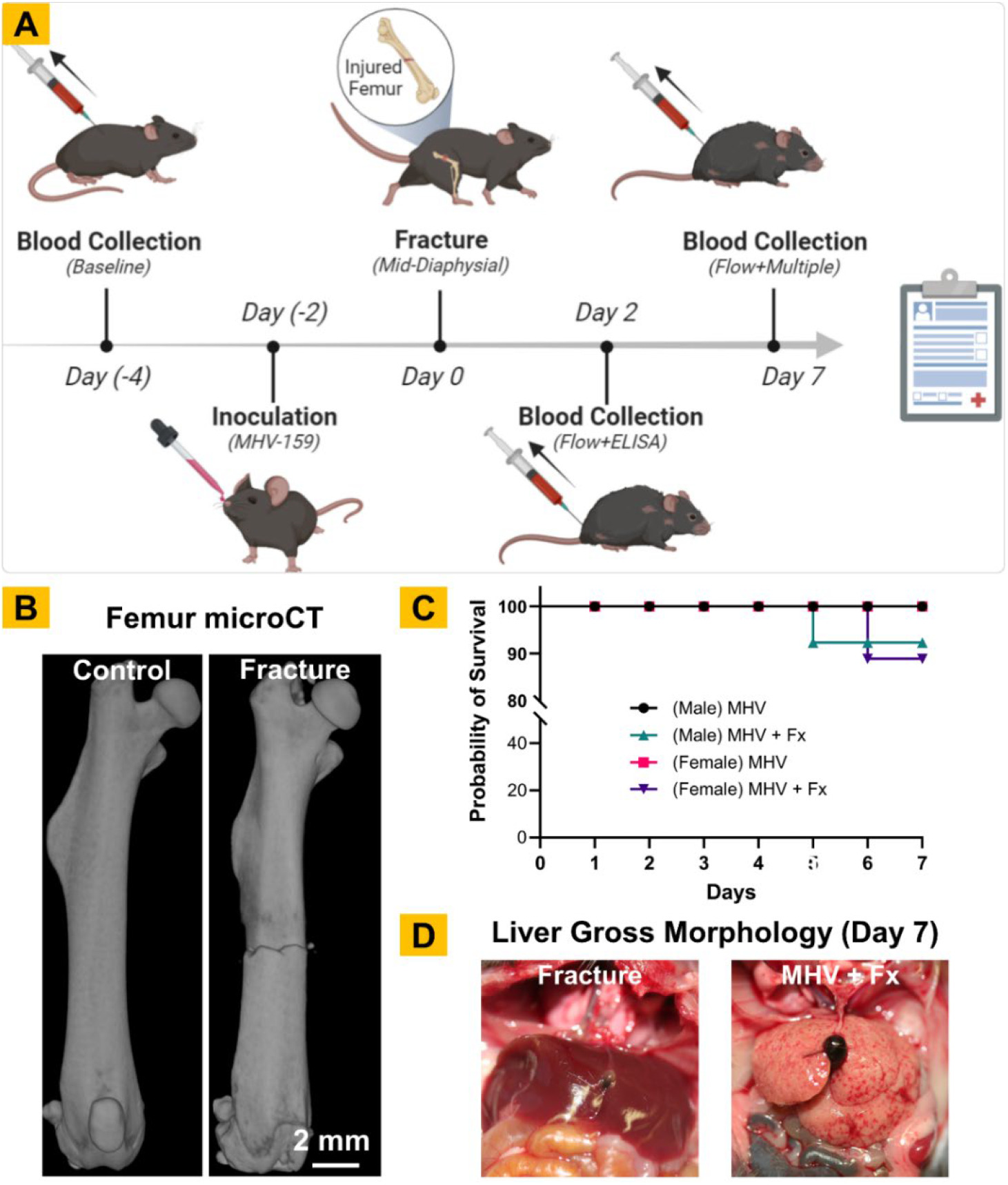
Schematic of the Study and Procedures. **A)** Timeline of the blood collection, inoculation, fracture, and endpoint for the study. **B)** Representative microCT images of control and fractured mouse femur on day 7, **C)** Kaplan– Meier curve showing percent survival in each group over a week post-fracture. **D)** Representative gross image of the liver collected on day 7 from uninfected and MHV-infected mice.

All mice in the control group remained healthy, active, and alert throughout the experiment. Mice in the FX group had reduced mobility for a few days post-fracture but showed signs of recovery within 2-3 days. In contrast, within 1-2 days post-inoculation, all infected mice (MHV-Sham and MHV+FX groups) displayed signs of illness, including hunched posture, wincing, shivering, and general lethargy. These symptoms persisted throughout the duration of the experiment.

In the MHV+FX group, the combination of infection and fracture led to a more severe decline, with a 20% mortality rate, compared to 0% in other groups (**Fig.1C**). Mice that survived were sacrificed 7 days post-fracture. At the time of death or sacrifice, the mice were dissected via thoracotomy for a visual inspection of organ health. Liver damage was more pronounced in the MHV+FX (**Fig.1D**), suggesting that the dual burden of infection and injury severely compromises health.

### 3.2. Fractures in MHV-Infected Mice Result in Systemic Immune Dysregulation

The dynamics of immune cell populations in the peripheral blood were quantified using a flow cytometry analysis of the whole blood, which showed significant differences between treatment groups and sexes **(Fig. 2 and 3)**.

**Figure 2.**
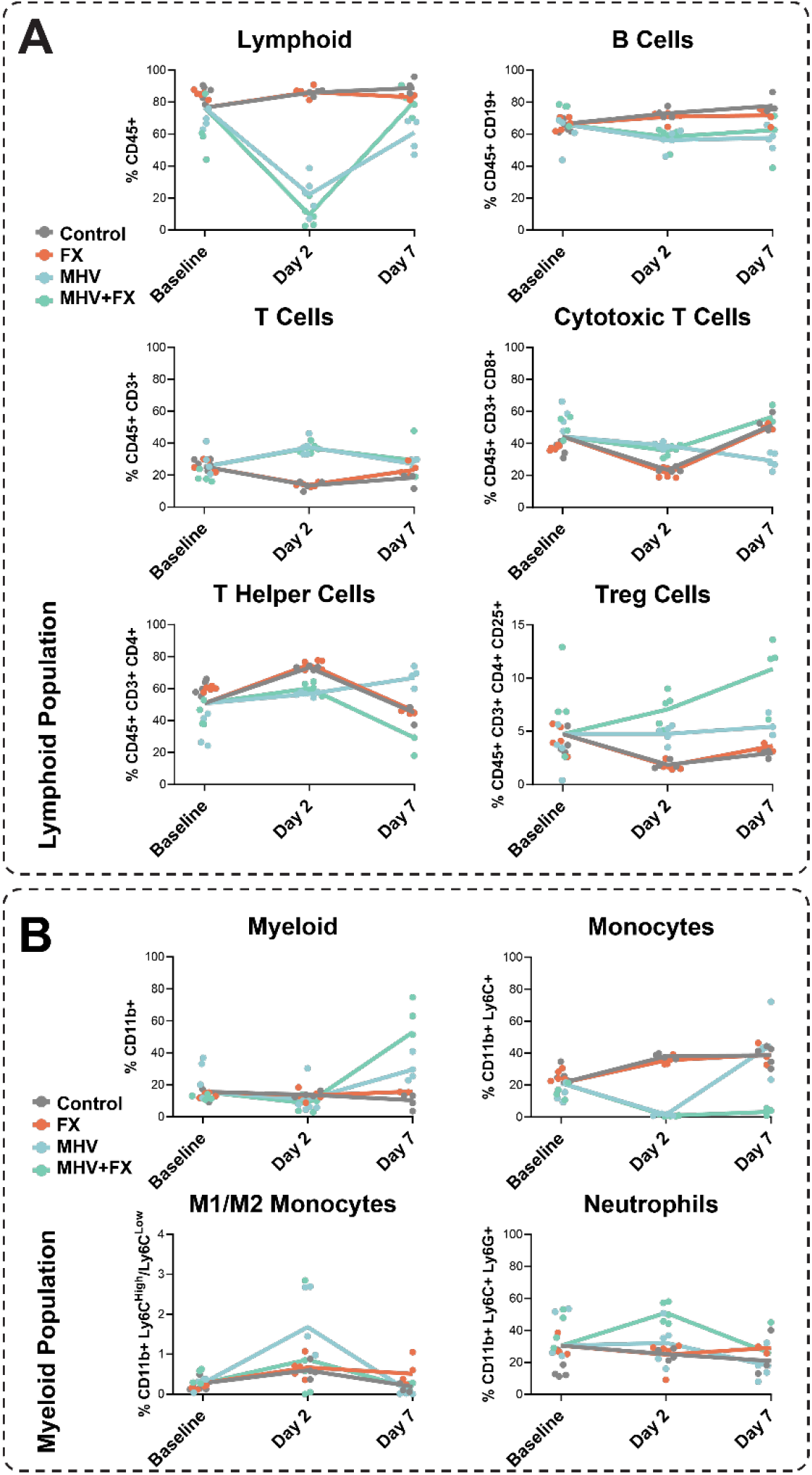
Immune Cell Populations of Female Mice Over Time. Immune cell profiles were assessed at Baseline, Day 2, and Day 7 post-fracture. **(A)** Lymphoid populations, including B cells, T cells, T helper cells, cytotoxic T cells, and T regulatory cells, showed temporal variation, with some populations increasing at Day 2 in the MHV+Fx group. **(B)** Myeloid populations, such as monocytes, neutrophils, macrophages (M1 and M2 subtypes), and dendritic cells, demonstrated dynamic changes, with neutrophils and M1 macrophages peaking at Day 2 in the MHV+Fx group. Values are expressed as individual data points with line plots representing group means. Treatment groups: Control (grey), Fx (orange), MHV (light blue), and MHV+Fx (light green).

**Figure 3.**
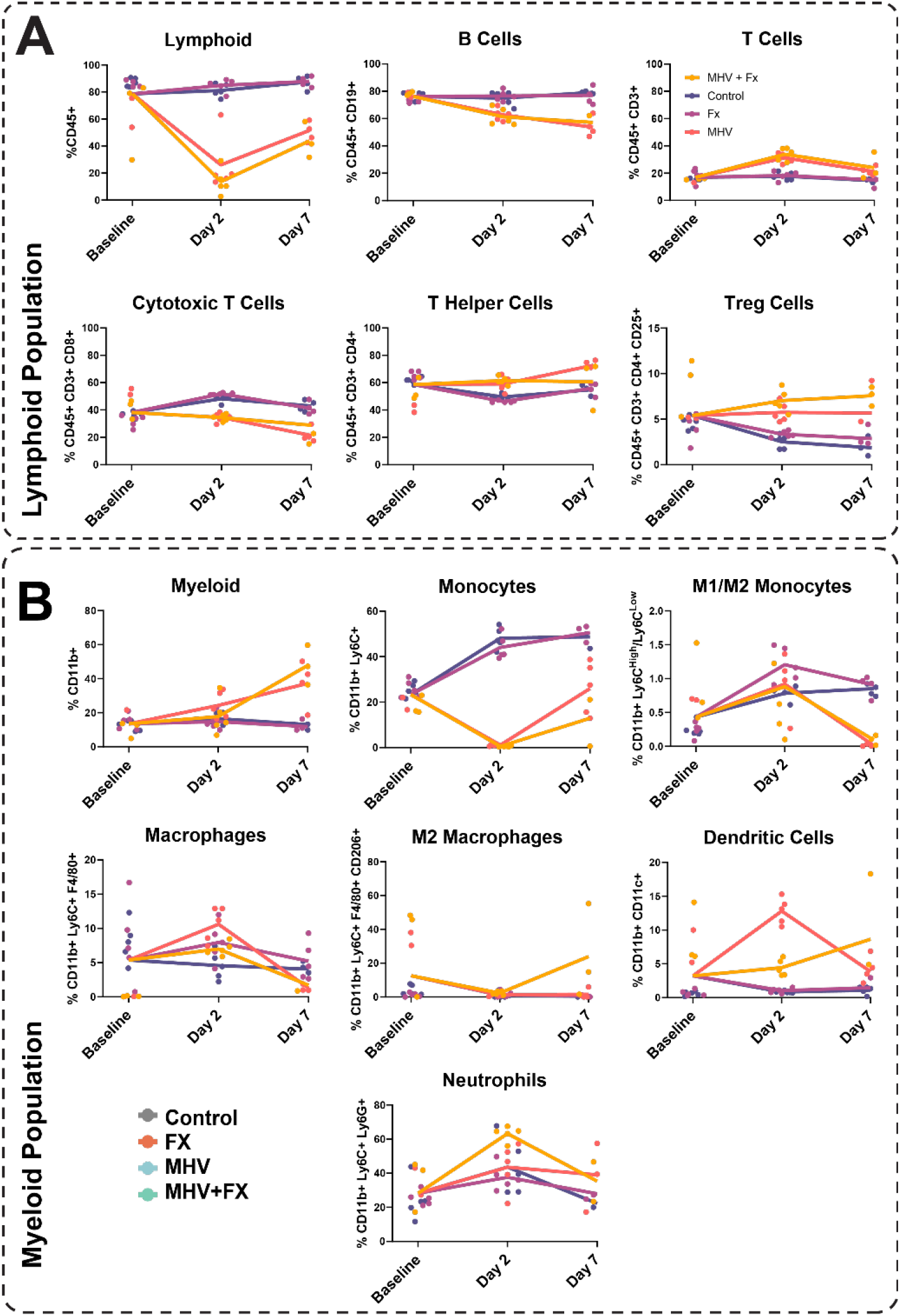
Immune Cell Populations of Male Mice Over Time. Immune cell profiles were assessed at Baseline, Day 2, and Day 7 post-fracture. **(A)** Lymphoid populations, including B cells, T cells, T helper cells, cytotoxic T cells, and T regulatory cells, displayed distinct temporal patterns, with certain subpopulations peaking at Day 2 in the MHV+Fx group. **(B)** Myeloid populations, such as monocytes, neutrophils, macrophages, and dendritic cells, exhibited varying trends, with neutrophils and macrophages showing elevated levels in the MHV+Fx group at Day 2. Values are expressed as individual data points with line plots representing group means. Treatment groups: Control (dark purple), Fx (light purple), MHV (pink), and MHV+Fx (yellow).

In female mice, MHV infection and the combined MHV+FX condition showed significant reductions in lymphoid cells, including B cells and T helper cells, as well as monocytes, when compared to the Control and FX groups. In contrast, CD8+ cytotoxic T cells and T regulatory cells (Tregs) showed a significant increase in the MHV and MHV+FX groups compared to the other treatment groups. However, no statistically significant differences were observed between the MHV and MHV+FX groups at this time point. By **Day 7**, the immune landscape continued to evolve. In the MHV group, there was a pronounced reduction in lymphoid and B cell populations compared to both the Control and FX groups, while T cells, T helper cells, and myeloid cells significantly increased. Interestingly, the cytotoxic T-cell population in the MHV group was significantly lower than in all other groups, indicating potential immune exhaustion or dysregulation in response to viral infection. In the MHV+FX group, the monocyte population was significantly reduced compared to all other treatment groups, suggesting impaired immune capacity in response to the combined stress of infection and trauma.

Several significant temporal shifts were also noted within the treatment conditions. Most notably, in the MHV group, the ratio of the M1/M2 monocytes (pro-inflammatory to anti-inflammatory phenotype) significantly increased from baseline, indicating a sustained pro-inflammatory state not seen in other groups. Additionally, in the MHV+FX group, the Treg population showed a significant elevation on both Days 2 and 7 compared to baseline, while other groups displayed either reduced or unchanged Treg levels over time. This sustained Treg activation in the MHV+FX group may indicate an ongoing attempt by the immune system to counterbalance the heightened inflammatory state but could also reflect immune suppression in the context of severe immune dysregulation.

Male mice mirrored the female immune response with a few key differences. On days 2 and 7, male mice in the MHV and MHV+FX groups demonstrated a decrease in cytotoxic T cells and an increase in T helper cells, contrasting the trend observed in female mice. This suggests that male and female mice may activate distinct immune pathways when facing both infection and injury. By day 7, the monocyte population in female mice had returned to levels similar to those in the Control and FX groups. However, in male mice, the monocyte population in the MHV+FX group remained significantly reduced. This persistent reduction may indicate a delayed or impaired immune response in males. Additionally, the M1/M2 monocyte ratio was significantly decreased at day 7 in both the MHV and MHV+FX groups for males, pointing to a shift towards a more anti-inflammatory or immunosuppressive state, which could impact healing and infection resolution. The dendritic cell population on day 2 was significantly elevated in the MHV and MHV+FX groups compared to the Control and FX groups, with the MHV group showing a greater increase than the MHV+FX group. This elevation was observed in both sexes but was more pronounced in males, suggesting heightened antigen-presenting activity in response to the viral infection. When examining changes over time within each condition, male responses closely paralleled those of females, with some notable differences. In particular, neutrophil levels in the MHV+FX group significantly increased compared to baseline and day 7, whereas neutrophils in other groups remained unchanged or decreased. This sustained neutrophil presence in males could contribute to prolonged inflammation, potentially affecting tissue repair and resolution of the injury (**Fig. 3**).

These findings suggest that both MHV infection and combined MHV+FX treatment conditions independently influence systemic cell populations in a sex-specific manner. A complete list of Holm-Sidak comparison p-values is provided in the **Supplementary Tables 1 - 4**.

### 3.3. Sex-specific Changes in Circulating Inflammatory Factors

Circulating inflammatory cytokines, chemokines, and growth factors were analyzed using a multiplex assay, which revealed notable sex-specific differences. **(Fig. 4 and 5)**.

**Figure 4.**
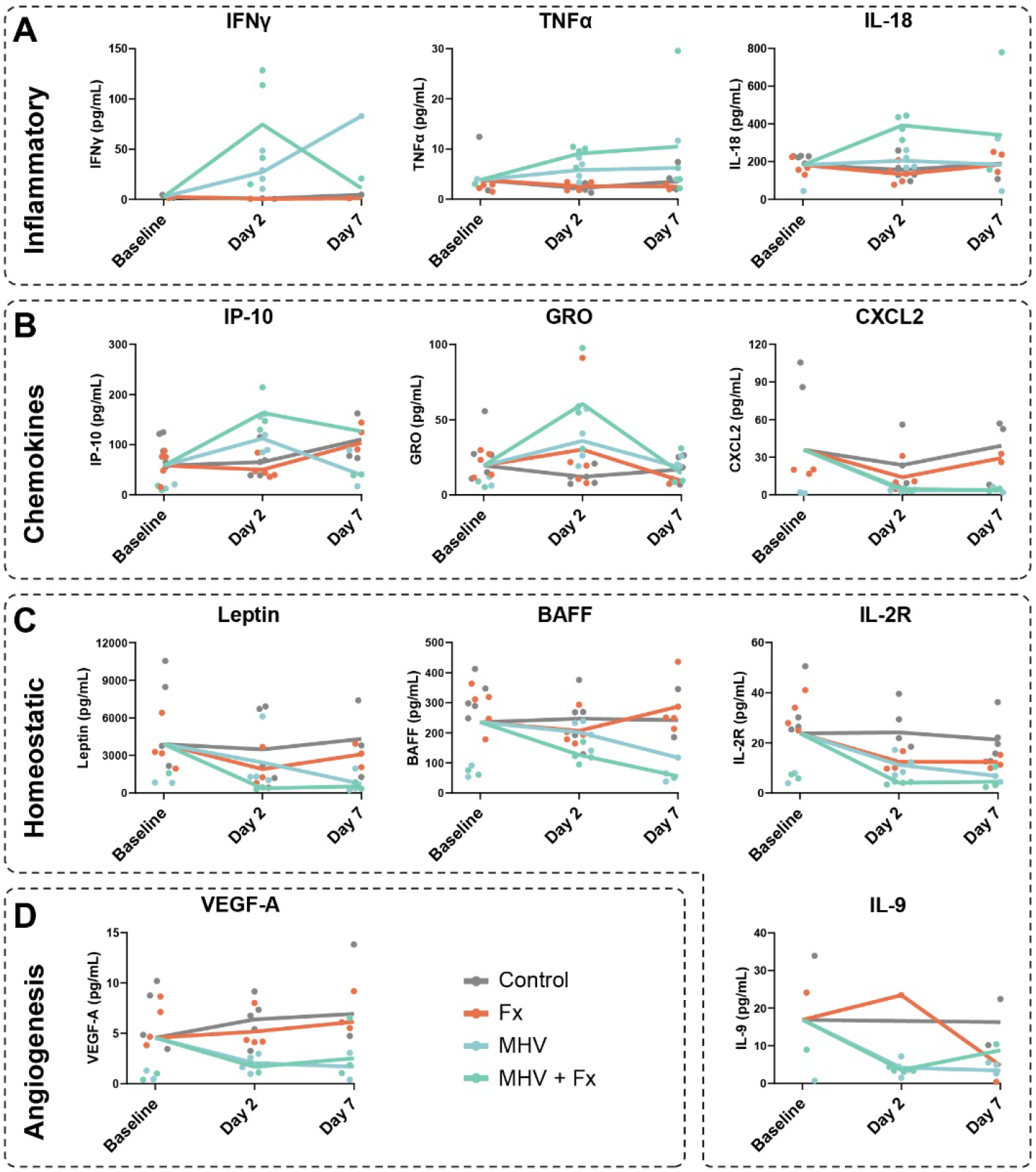
Temporal changes in cytokine levels in Blood Plasma of Female mice across inflammatory, chemokine, homeostatic, and angiogenic categories in response to experimental conditions. Cytokine and chemokine profiles at Baseline, Day 2, and Day 7 after treatment. **(A)** Pro-inflammatory cytokines IFN-γ, TNF-α, and IL-18 showed significant increases at Day 2 in the MHV + Fx group compared to controls. **(B)** Chemokines IP-10 and GRO peaked at Day 2, while CXCL2 showed minimal changes. **(C)** Homeostatic and regulatory cytokines, including Leptin, BAFF, IL-2R, and IL-9, demonstrated declining trends over the 7-day period. **(D)** VEGF-A levels declined over time, suggesting reduced angiogenic activity. Values are expressed as individual data points with line plots representing group means. Treatment groups: Control (grey), Fx (orange), MHV (light blue), and MHV+Fx (light green).

**Figure 5.**
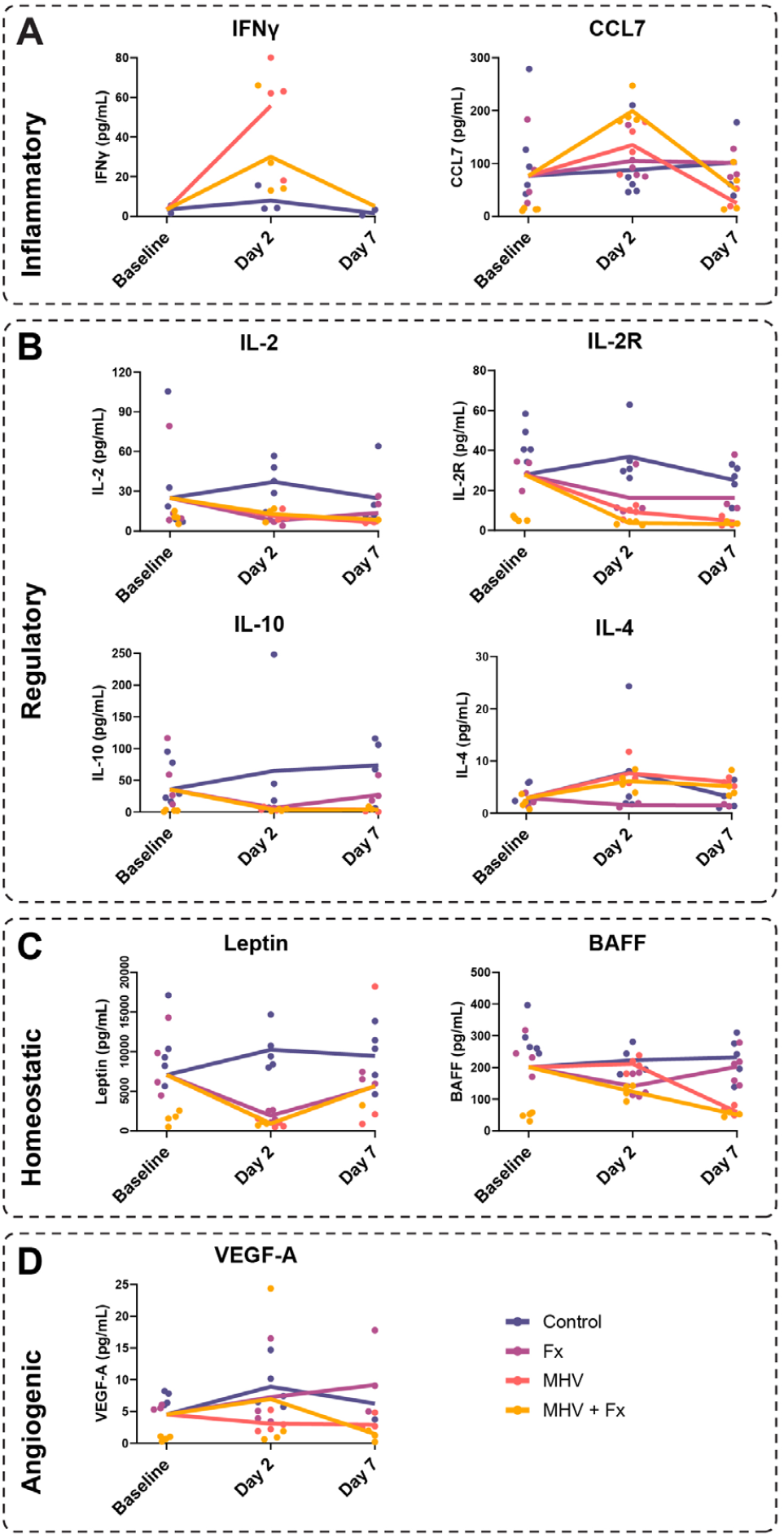
Temporal changes in cytokine levels in Blood Plasma of Male Mice across inflammatory, regulatory, homeostatic, and angiogenic functional groups in response to experimental conditions. **(A)** Pro-inflammatory cytokines (IFN-γ, CCL7) exhibited a transient increase on Day 2, particularly in the MHV and MHV+Fx groups, suggesting an acute immune response. **(B)** Regulatory cytokines (IL-2, IL-2R, IL-10, IL-4) displayed varied dynamics, with IL-2 peaking at Day 2 in the MHV+Fx group, while IL-2R, IL-10, and IL-4 remained relatively stable across time points. **(C)** Homeostatic factors (Leptin, BAFF) showed a marked reduction by Day 7 in MHV and MHV+Fx groups, indicating potential metabolic shifts or systemic effects. **(D)** Angiogenic factor VEGF-A exhibited a transient increase on Day 2 in the MHV+Fx group, suggesting vascular involvement in the response to treatments. Values are expressed as individual data points with line plots representing group means. Treatment groups: Control (dark purple), Fx (light purple), MHV (pink), and MHV+Fx (yellow).

In female mice, IFNγ was detected exclusively in virally infected mice at day 2, confirming successful inoculation and infection. Of the 48 immune factors tested, only those represented in **Figure 4** showed significant changes (full list in the **supplementary** material). On day 2, IL-18 exhibited a significant 2-fold increase in the MHV+FX group compared to all other groups. TNFα and IP-10 were significantly elevated in the MHV+FX group compared to the Control and FX groups but not the MHV group. VEGF-A remained significantly reduced in the MHV group compared to the FX group. Additionally, Leptin decreased significantly in both the MHV and MHV+FX groups, while IL-2R levels in the MHV group significantly decreased from day 2 to day 7.

In male mice, as in females, IFNγ was detected only in virally infected groups at day 2, confirming infection. Of the 48 immune factors tested in male mice, only those represented in **Figure 5** were notable (full results in supplementary material).

On **day 2**, IL-2R significantly decreased in the MHV+FX group compared to the Control, with a further decrease by day 7 in both MHV and MHV+FX groups compared to Control. Unlike in females, Leptin levels were significantly reduced at day 2 across the FX, MHV, and MHV+FX groups, with no significant change by day 7. BAFF levels decreased in the FX and MHV groups at day 2 but remained unchanged in the MHV+FX group. By **day 7**, BAFF decreased in the MHV+FX group compared to the Control and FX groups, consistent with findings in females. CCL7 was notably increased in the MHV+FX group compared to the FX group at day 2, and by day 7, both BAFF and CCL7 levels had decreased significantly in the MHV+FX group compared to day 2. Finally, at day 7 IL-4 significantly increased only in the MHV group compared to the FX group, with IL-4 levels also increasing from baseline to day 2 in the MHV group.

These findings underscore that both MHV infection and the combined MHV+FX condition elicit distinct immune responses, with significant sex-specific variations. Female and male mice shared common alterations in only four immune factors: IFNγ, Leptin, BAFF, and VEGF-A. However, male mice exhibited unique patterns in Leptin, BAFF, CCL7, and IL-4 levels that were not observed in females. The sex-specific immune modulation highlights the importance of considering gender in immune response studies.

### 3.4. Statistical Analysis Reveals Sex-Specific Differences and Compounding Effects of Infection and Fracture

Principal Component Analysis (PCA) was conducted to simplify the complex datasets illustrated in **Figures 2** through **5** and to identify the primary patterns of variation within the data. The resulting visualizations (**Fig. 6A–C**) elucidate how fracture (FX), viral infection (MHV), and their combination (MHV+FX) influence immune responses

**Figure 6:**
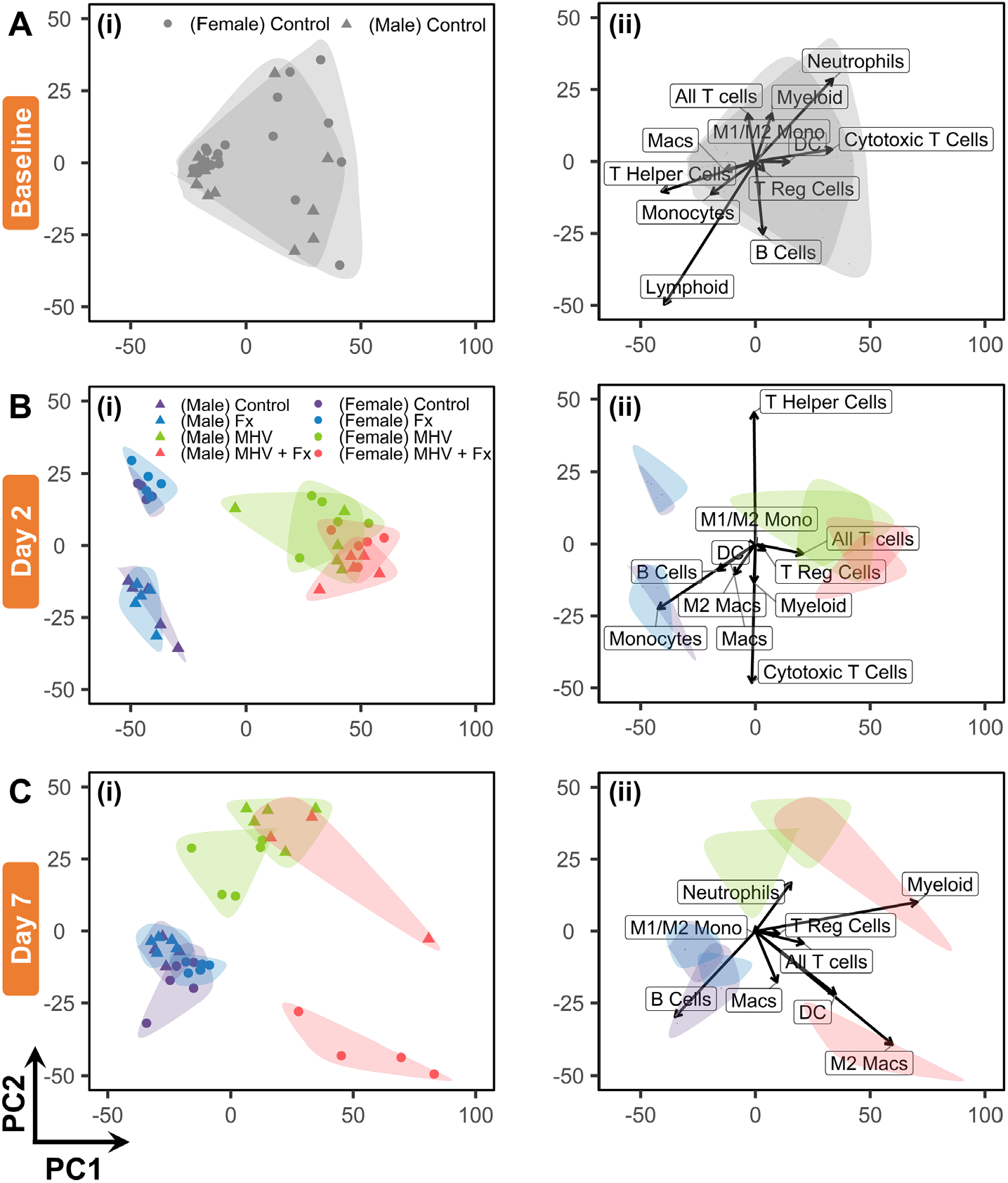
Principal Component Analysis (PCA) of Experimental and Control Groups. PCA plots show principal component scores (PC1 and PC2) across experimental groups, highlighting variance and treatment effects. **(A)** Baseline Immune Profiles: (i) Score plot showing minimal differences between male and female cohorts at baseline. (ii) Loading map identifying cytotoxic T cells, B cells, and M2 macrophages as key contributors to variance. **(B)** Day 2 Post-Treatment: (i) Overlap between control and fracture-only (FX) groups, with distinct separation of MHV and MHV+FX groups due to immune modulation. (ii) T cells, dendritic cells, and M2 macrophages drive observed variance. **(C)** Day 7 Post-Treatment: (i) MHV+FX group, particularly females, shows distinct clustering, reflecting compounded immune responses. (ii) M2 macrophages are key drivers of variance in the MHV+FX group. Each dot represents an individual mouse, color-coded by experimental group.

**Figure 6A(i)** shows the baseline values of immune cell populations in the blood which show no significant difference between male and female cohorts at the outset. The baseline PCA clusters for both sexes are virtually identical. Figure **6A(ii)** illustrates the corresponding PCA loading map, highlighting the major variables driving the dataset’s variance.

By day 2, sex-specific differences become evident (**Fig. 6B(i)**). PCA reveals substantial overlap between control and fracture-only (FX) groups in both sexes, suggesting that fractures alone minimally influence immune cell populations and soluble factors. However, exposure to MHV significantly alters immune responses, with distinct clustering of MHV and MHV+FX groups compared to controls. These distinctions are largely driven by T cell populations, as evidenced in the loading maps (**Fig. 6B(ii)**).

By day 7, pronounced differences emerge in the MHV+FX group, particularly in female mice (**Fig. 6C(i)**). This group is distinctly separated along both principal components (PC1 and PC2), suggesting a unique compounded response to simultaneous infection and fracture. The corresponding PCA loading plot (**Fig. 6C(ii)**) attributes this separation primarily to changes in M2 macrophage populations, underscoring their significant contribution to the immune response under dual challenge conditions.

These findings highlight the necessity of accounting for both sex-specific differences and the interaction between infection and injury to fully understand the immune responses elicited under these conditions. The loadings plot reveals key insights into how specific immune cell populations modulate the immune landscape, emphasizing the distinct roles of T cell subtypes and macrophages in mounting the immune responses under varying conditions.

## 4. DISCUSSION

Dysregulated inflammation is a key disruptor of healing in musculoskeletal injuries, often hindering the body’s ability to recover effectively. Coronavirus infections, such as those caused by SARS-CoV-2, are notorious for inducing a heightened systemic inflammatory response^22^, which can be further amplified by the inflammatory cascade triggered by fractures and other musculoskeletal traumas. The convergence of these inflammatory events creates a perfect storm, complicating patient recovery and driving up mortality rates^23^. Our results provide compelling evidence of the compounding effects of viral infection and trauma on mortality and immune dysregulation in a murine model, along with significant sex-specific differences in immune response. Our results indicate that the combination of MHV infection and fracture markedly worsens clinical outcomes, emphasizing the critical need to understand the intertwined effects of these two stressors on the immune system. Below, we discuss the two main phenomena observed: the compounding effect of infections and trauma and sex-specific differences in immune responses.

### 4.1. Compounding Effects of Infection and Trauma

Our findings demonstrate a significant increase in mortality in mice subjected to both MHV infection and fracture (MHV+FX), with a 20% mortality rate in the MHV+FX group compared to 0% in all other groups. This synergistic effect of viral infection and trauma suggests a profound disruption of homeostatic immune responses, leading to exacerbated systemic inflammation and organ damage, as evidenced by severe liver damage in the MHV+FX group. The observed compounding effects can be hypothesized to result from an overwhelming inflammatory response triggered by the simultaneous presence of viral infection and musculoskeletal trauma. Viral infections are known to initiate a robust immune response characterized by the activation of innate immune cells, including macrophages, neutrophils, and dendritic cells, and the production of pro-inflammatory cytokines such as IFNγ and TNFα^24^. In parallel, trauma induces a distinct but overlapping inflammatory cascade, which includes the release of damage-associated molecular patterns (DAMPs) that further amplify the immune response^25^. The convergence of these pathways may lead to a “cytokine storm,” characterized by excessive and dysregulated cytokine production, as observed in our study, where IFNγ, IL-18 and TNFα were significantly elevated in the MHV+FX group. This heightened inflammatory state likely contributes to increased tissue damage, organ dysfunction, and mortality.

The reduction in circulating lymphoid and myeloid cell populations, particularly B cells and monocytes, in MHV and MHV+FX groups supports this hypothesis. Viral infections often drive lymphocyte apoptosis or redistribution to tissues, and trauma can exacerbate this effect, resulting in immune-paralysis and increased susceptibility to secondary infections^26^. Moreover, the significant reduction in monocyte populations in the MHV+FX group suggests a depletion of these critical immune cells, which may impair the body’s ability to clear infection and repair tissue, further compounding the adverse outcomes observed.

Interestingly, the T regulatory cell (Treg) population was significantly elevated in the MHV+FX group compared to baseline, which may represent a compensatory mechanism to counteract the excessive inflammation. However, while Tregs play a role in resolving inflammation, their overactivation in the context of severe infection and trauma can paradoxically suppress effective immune responses, contributing to a state of immune exhaustion^27^. This immune dysregulation underscores the importance of understanding the delicate balance between immune activation and suppression in complex clinical scenarios involving infection and injury.

### 4.2. Sex-Specific Differences in Immune Response

Our data reveal significant sex-specific differences in immune responses to MHV infection and fracture. Female mice exhibited a distinct pattern of immune cell population dynamics and inflammatory cytokine expression compared to males. For instance, the MHV+FX group of females showed a significant increase in Treg cells and a pronounced shift in the M1/M2 monocyte ratio, indicating a sex-specific polarization of immune responses. These findings align with growing evidence that sex hormones and genetic differences influence immune function^28^.

In contrast, male mice in the MHV and MHV+FX groups exhibited a persistent reduction in cytotoxic T cells and an increase in T helper (Th) cells, opposite to the trends observed in females. This suggests that male and female mice utilize distinct immunological strategies to cope with the combined stress of infection and trauma. Males appeared to favor a Th2-skewed response, which is typically associated with antibody production and wound healing but may be less effective in controlling viral infections^29^. Conversely, females showed a more pronounced Th1 and Treg response, which, while effective against pathogens, could exacerbate inflammation when dysregulated.

The PCA analysis further highlights the impact of sex on the immune landscape, with distinct separations observed between male and female mice in the MHV+FX group. Notably, cytotoxic T cells, M2 macrophages, and dendritic cells contributed significantly to the observed variance, emphasizing their critical roles in mediating the immune response under different conditions. Furthermore, females showed a significant increase in circulating IL-18 and TNFα levels in the MHV+FX group, which was not mirrored in males. This suggests that females may mount a more exaggerated inflammatory response to the combined stressors, potentially explaining the observed differences in immune cell dynamics and outcomes. Similar patterns are seen clinically, with females displaying stronger inflammatory responses to COVID-19 than males^30^, though this heightened response may increase the risk of cytokine storm in the presence of comorbidities^7^.

## 5. IMPLICATIONS AND FUTURE DIRECTIONS

The current findings have important implications for understanding the interplay between infection, trauma, and immune response in a sex-specific context. The observed compounding effects of MHV infection and fracture highlight the need for integrated therapeutic approaches that address both infection control and immune modulation. Moreover, the pronounced sex-specific differences emphasize the importance of personalized treatment strategies that consider the patient’s sex, particularly in the management of complex conditions involving both infection and physical trauma.

Despite its significance, the study has certain limitations. The survival rate analysis was conducted over a relatively short time frame, and biomechanical assessments of bone healing—typically performed at three weeks in murine models—were not included. Addressing these gaps in future studies will provide a more comprehensive understanding of the long-term effects of these comorbid conditions.

Future research should aim to elucidate the molecular mechanisms driving the observed sex-specific immune responses, including the roles of sex hormones, genetic factors, and microbiome differences. Understanding these mechanisms could lead to the development of targeted therapies that optimize immune responses and improve outcomes in patients experiencing the dual burden of infection and trauma.

## Reference

1. Dong, E., Du, H. & Gardner, L. An interactive web-based dashboard to track COVID-19 in real time. The Lancet infectious diseases 20, 533–534 (2020).

2. Seighali, N. et al. The global prevalence of depression, anxiety, and sleep disorder among patients coping with Post COVID-19 syndrome (long COVID): a systematic review and meta-analysis. BMC psychiatry 24, 105 (2024).

3. Siddiqi, H.K. & Mehra, M.R. COVID-19 illness in native and immunosuppressed states: A clinical– therapeutic staging proposal. The journal of heart and lung transplantation 39, 405–407 (2020).

4. Mehta, P. et al. COVID-19: consider cytokine storm syndromes and immunosuppression. The lancet 395, 1033–1034 (2020).

5. Ruan, Q., Yang, K., Wang, W., Jiang, L. & Song, J. Clinical predictors of mortality due to COVID-19 based on an analysis of data of 150 patients from Wuhan, China. Intensive care medicine 46, 846–848 (2020).

6. Takahashi, T. et al. Sex differences in immune responses that underlie COVID-19 disease outcomes. Nature 588, 315–320 (2020).

7. Anca, P.S., Toth, P.P., Kempler, P. & Rizzo, M. Gender differences in the battle against COVID-19: Impact of genetics, comorbidities, inflammation and lifestyle on differences in outcomes. International journal of clinical practice 75 (2021).

8. Klein, S.L. & Flanagan, K.L. Sex differences in immune responses. Nature Reviews Immunology 16, 626–638 (2016).

9. Collins, M.K., McCutcheon, C.R. & Petroff, M.G. Impact of Estrogen and Progesterone on Immune Cells and Host–Pathogen Interactions in the Lower Female Reproductive Tract. The Journal of Immunology 209, 1437–1449 (2022).

10. Furman, D. et al. Systems analysis of sex differences reveals an immunosuppressive role for testosterone in the response to influenza vaccination. Proceedings of the National Academy of Sciences 111, 869–874 (2014).

11. Scully, E.P., Haverfield, J., Ursin, R.L., Tannenbaum, C. & Klein, S.L. Considering how biological sex impacts immune responses and COVID-19 outcomes. Nat Rev Immunol 20, 442–447 (2020).

12. Haffner-Luntzer, M., Fischer, V. & Ignatius, A. Differences in Fracture Healing Between Female and Male C57BL/6J Mice. Frontiers in physiology 12 (2021).

13. Fischer, V. et al. Influence of Menopause on Inflammatory Cytokines during Murine and Human Bone Fracture Healing. Int J Mol Sci 19 (2018).

14. Drucker, D.J. Coronavirus infections and type 2 diabetes—shared pathways with therapeutic implications. Endocrine reviews 41, bnaa011 (2020).

15. Bornstein, S.R., Dalan, R., Hopkins, D., Mingrone, G. & Boehm, B.O. Endocrine and metabolic link to coronavirus infection. Nature Reviews Endocrinology 16, 297–298 (2020).

16. Mi, B. et al. Characteristics and early prognosis of COVID-19 infection in fracture patients. The Journal of Bone and Joint Surgery. American Volume 102, 750 (2020).

17. Vives, J.M.M. et al. Mortality rates of patients with proximal femoral fracture in a worldwide pandemic: preliminary results of the Spanish HIP-COVID observational study. The Journal of bone and joint surgery. American volume (2020).

18. Campolina-Silva, G. et al. Dietary vitamin D mitigates Coronavirus-Induced Lung inflammation and damage in mice. Viruses 15, 2434 (2023).

19. Arévalo, A. et al. Ivermectin reduces in vivo coronavirus infection in a mouse experimental model. Scientific reports 11, 7132 (2021).

20. Bonnarens, F. & Einhorn, T.A. Production of a standard closed fracture in laboratory animal bone. Journal of orthopaedic research 2, 97–101 (1984).

21. Bartnikowski, M., Bartnikowski, N., Woloszyk, A., Matthys, R. & Glatt, V. Genetic variation in mice affects closed femoral fracture pattern outcomes. Injury 50, 639–647 (2019).

22. Buicu, A.L., Cernea, S., Benedek, I., Buicu, C.F. & Benedek, T. Systemic Inflammation and COVID-19 Mortality in Patients with Major Noncommunicable Diseases: Chronic Coronary Syndromes, Diabetes and Obesity. J Clin Med 10 (2021).

23. Jagadeesh, N. et al. COVID-19 infection increases mortality and complications in patients with neck of femur fracture. Cureus 14 (2022).

24. Takeuchi, O. & Akira, S. Pattern Recognition Receptors and Inflammation. Cell 140, 805–820 (2010).

25. Relja, B. & Land, W.G. Damage-associated molecular patterns in trauma. European Journal of Trauma and Emergency Surgery 46, 751–775 (2020).

26. Hotchkiss, R.S., Monneret, G. & Payen, D. Sepsis-induced immunosuppression: from cellular dysfunctions to immunotherapy. Nature Reviews Immunology 13, 862–874 (2013).

27. Romano, M., Tung, S.L., Smyth, L.A. & Lombardi, G. Treg therapy in transplantation: a general overview. Transplant International 30, 745–753 (2017).

28. Klein, S.L. & Flanagan, K.L. Sex differences in immune responses. Nat Rev Immunol 16, 626–638 (2016).

29. vom Steeg, L.G. & Klein, S.L. SeXX Matters in Infectious Disease Pathogenesis. PLOS Pathogens 12, e1005374 (2016).

30. Conti, P. & Younes, A. Coronavirus COV-19/SARS-CoV-2 affects women less than men: clinical response to viral infection. J Biol Regul Homeost Agents 34, 339–343 (2020).

