## Supplemental Material for "Sexual Dimorphism in Systemic Inflammatory Responses to Femur Fracture in Mice Infected with SARS-CoV-2-Like Virus"

### Supplementary Material

**Supplementary Table 1:** Female Flow Cytometry Holm-Sidak Pairwise Comparison (Condition within Time Point) (p-value)

| | | Lymphoid | B cells | T cells | CD8+ T cells | CD4+ T cells | CD25+ T cells | Myeloid | Monocytes | M1/M2 Monocytes | Neutrophils |
|---|---|---|---|---|---|---|---|---|---|---|---|
| BaseLine | Control vs. Fracture | >0.9999 | >0.9999 | >0.9999 | >0.9999 | >0.9999 | >0.9999 | >0.9999 | >0.9999 | >0.9999 | >0.9999 |
| | Control vs. MHV | >0.9999 | >0.9999 | >0.9999 | >0.9999 | >0.9999 | >0.9999 | >0.9999 | >0.9999 | >0.9999 | >0.9999 |
| | Control vs. MHV + Fracture | >0.9999 | >0.9999 | >0.9999 | >0.9999 | >0.9999 | >0.9999 | >0.9999 | >0.9999 | >0.9999 | >0.9999 |
| | Fracture vs. MHV | >0.9999 | >0.9999 | >0.9999 | >0.9999 | >0.9999 | >0.9999 | >0.9999 | >0.9999 | >0.9999 | >0.9999 |
| | Fracture vs. MHV + Fracture | >0.9999 | >0.9999 | >0.9999 | >0.9999 | >0.9999 | >0.9999 | >0.9999 | >0.9999 | >0.9999 | >0.9999 |
| | MHV vs. MHV + Fracture | >0.9999 | >0.9999 | >0.9999 | >0.9999 | >0.9999 | >0.9999 | >0.9999 | >0.9999 | >0.9999 | >0.9999 |
| Day 2 | Control vs. Fracture | 0.8248 | 0.6219 | 0.9694 | 0.2439 | 0.2584 | 0.6662 | 0.9702 | 0.159 | 0.9524 | 0.9489 |
| | Control vs. MHV | 0.0007* | 0.0106* | 0.0004* | <0.0001* | <0.0001* | 0.002* | 0.9702 | <0.0001* | 0.3338 | 0.6871 |
| | Control vs. MHV + Fracture | <0.0001* | 0.016* | <0.0001* | 0.0021* | 0.0022* | 0.0049* | 0.343 | <0.0001* | 0.9524 | 0.0011* |
| | Fracture vs. MHV | 0.0006* | 0.016* | 0.0012* | 0.0001* | 0.0001* | 0.0018* | 0.9702 | <0.0001* | 0.3367 | 0.6871 |
| | Fracture vs. MHV + Fracture | <0.0001* | 0.0212* | 0.0001* | 0.0007* | 0.0006* | 0.0049* | 0.4186 | <0.0001* | 0.9524 | 0.0048* |
| | MHV vs. MHV + Fracture | 0.1651 | 0.6219 | 0.9694 | 0.2439 | 0.1872 | 0.0568 | 0.9702 | 0.159 | 0.7092 | 0.1233 |
| Day 7 | Control vs. Fracture | 0.1093 | 0.2839 | 0.3306 | 0.7674 | 0.6723 | 0.1684 | 0.1307 | 0.9097 | 0.2765 | 0.6165 |
| | Control vs. MHV | 0.0118* | 0.0014* | 0.0477* | 0.0005* | 0.0012* | 0.0598 | 0.0082* | 0.8536 | 0.1023 | 0.9418 |
| | Control vs. MHV + Fracture | 0.1017 | 0.427 | 0.5121 | 0.3003 | 0.079 | 0.0801 | 0.1179 | 0.0003* | 0.5944 | 0.9418 |
| | Fracture vs. MHV | 0.0406 | 0.0097* | 0.5121 | 0.0011* | 0.0034* | 0.0963 | 0.0483* | 0.8536 | 0.142 | 0.2298 |
| | Fracture vs. MHV + Fracture | 0.1093 | 0.5885 | 0.6863 | 0.251 | 0.079 | 0.0801 | 0.1307 | 0.0003* | 0.2618 | 0.9418 |
| | MHV vs. MHV + Fracture | 0.1614 | 0.6105 | 0.7544 | 0.0005* | 0.0034* | 0.0963 | 0.1307 | 0.0221* | 0.3527 | 0.9107 |

**Supplementary Table 2:** Female Flow Cytometry Holm-Sidak Pairwise Comparison (Time Point within Condition) (p-value)

| | | Lymphoid | B cells | T cells | CD8+ T cells | CD4+ T cells | CD25+ T cells | Myeloid | Monocytes | M1/M2 Monocytes | Neutrophils |
|---|---|---|---|---|---|---|---|---|---|---|---|
| Control | Baseline vs Day 2 | 0.218 | 0.0292* | 0.0035* | 0.0275* | 0.0377* | 0.0268* | 0.4449 | 0.0383* | 0.0087* | 0.7032 |
| | Baseline vs Day 7 | 0.1559 | 0.0015* | 0.0071* | 0.1128 | 0.2487 | 0.0282* | 0.3463 | 0.0383* | 0.3789 | 0.5938 |
| | Day 2 vs Day 7 | 0.3195 | 0.0504 | 0.029* | 0.0089* | 0.0054* | 0.0282* | 0.4449 | 0.8711 | 0.0713 | 0.7032 |
| Fracture | Baseline vs Day 2 | 0.2163 | 0.2613 | 0.003* | 0.009* | 0.0131* | 0.0144* | 0.4621 | 0.0254* | 0.1128 | 0.7422 |
| | Baseline vs Day 7 | 0.2163 | 0.2613 | 0.4315 | 0.1416 | 0.3316 | 0.0754 | 0.9568 | 0.0254* | 0.354 | 0.8387 |
| | Day 2 vs Day 7 | 0.2163 | 0.3914 | 0.0056* | <0.0001* | <0.0001* | 0.0234* | 0.1597 | 0.3234 | 0.4915 | 0.7422 |
| MHV | Baseline vs Day 2 | 0.0041* | 0.0835 | 0.0248* | 0.1549 | 0.2259 | 0.9541 | 0.2774 | 0.0063* | 0.0449* | 0.8143 |
| | Baseline vs Day 7 | 0.0699 | 0.0835 | 0.4965 | 0.0191* | 0.0352* | 0.6626 | 0.0379* | 0.0112* | 0.0449* | 0.3191 |
| | Day 2 vs Day 7 | 0.0073* | 0.7526 | 0.0316* | 0.0159* | 0.0196* | 0.4131 | 0.065* | 0.0112* | 0.0449* | 0.3191 |
| MHV + Fracture | Baseline vs Day 2 | 0.0004* | 0.0718 | 0.0051* | 0.1437 | 0.1068 | 0.0383* | 0.0694 | 0.0047* | 0.4424 | 0.1176 |
| | Baseline vs Day 7 | 0.0457* | 0.8112 | 0.5722 | 0.1437 | 0.0972 | 0.0444* | 0.0529 | 0.0194* | 0.3733 | 0.5838 |
| | Day 2 vs Day 7 | 0.066* | 0.8112 | 0.5722 | 0.002* | 0.0052* | 0.1963 | 0.0694 | 0.1042 | 0.4424 | 0.1602 |

**Supplementary Table 3:** Male Flow Cytometry Holm-Sidak Pairwise Comparison (Condition within Time Point) (p-value)

| | | Lymphoid | B cells | T cells | CD8+ T cells | CD4+ T cells | CD25+ T cells | Myeloid | Monocytes | M1/M2 Monocytes | Macrophages | M2 Macrophages | Dendritic cells | Neutrophils |
|---|---|---|---|---|---|---|---|---|---|---|---|---|---|---|
| Baseline | Control vs. Fracture | >0.9999 | >0.9999 | >0.9999 | >0.9999 | >0.9999 | >0.9999 | >0.9999 | >0.9999 | >0.9999 | >0.9999 | >0.9999 | >0.9999 | >0.9999 |
| | Control vs. MHV | >0.9999 | >0.9999 | >0.9999 | >0.9999 | >0.9999 | >0.9999 | >0.9999 | >0.9999 | >0.9999 | >0.9999 | >0.9999 | >0.9999 | >0.9999 |
| | Control vs. MHV + Fracture | >0.9999 | >0.9999 | >0.9999 | >0.9999 | >0.9999 | >0.9999 | >0.9999 | >0.9999 | >0.9999 | >0.9999 | >0.9999 | >0.9999 | >0.9999 |
| | Fracture vs. MHV | >0.9999 | >0.9999 | >0.9999 | >0.9999 | >0.9999 | >0.9999 | >0.9999 | >0.9999 | >0.9999 | >0.9999 | >0.9999 | >0.9999 | >0.9999 |
| | Fracture vs. MHV + Fracture | >0.9999 | >0.9999 | >0.9999 | >0.9999 | >0.9999 | >0.9999 | >0.9999 | >0.9999 | >0.9999 | >0.9999 | >0.9999 | >0.9999 | >0.9999 |
| | MHV vs. MHV + Fracture | >0.9999 | >0.9999 | >0.9999 | >0.9999 | >0.9999 | >0.9999 | >0.9999 | >0.9999 | >0.9999 | >0.9999 | >0.9999 | >0.9999 | >0.9999 |
| Day 2 | Control vs. Fracture | 0.4367 | 0.841 | 0.7946 | 0.0746 | 0.0325 | 0.1339 | 0.7406 | 0.2336 | 0.1066 | 0.2105 | 0.9661 | 0.3672 | 0.8127 |
| | Control vs. MHV | 0.0099 | 0.018 | 0.0011 | 0.0005 | 0.0325 | 0.004 | 0.6568 | 0.0001 | 0.8962 | 0.0201 | 0.9661 | 0.0006 | 0.9968 |
| | Control vs. MHV + Fracture | <0.0001 | 0.0198 | 0.0019 | 0.0002 | 0.0007 | 0.0017 | 0.9182 | 0.0001 | 0.9621 | 0.2105 | 0.7006 | 0.0069 | 0.2086 |
| | Fracture vs. MHV | 0.0099 | 0.0057 | 0.0015 | 0.0007 | 0.024 | 0.0163 | 0.3073 | 0.0002 | 0.7279 | 0.3013 | 0.9661 | 0.0006 | 0.8127 |
| | Fracture vs. MHV + Fracture | <0.0001 | 0.0157 | 0.0019 | 0.0002 | 0.0008 | 0.0076 | 0.835 | 0.0002 | 0.8962 | 0.5155 | 0.8835 | 0.0069 | 0.0028 |
| | MHV vs. MHV + Fracture | 0.4367 | 0.841 | 0.5788 | 0.9755 | 0.395 | 0.1339 | 0.7406 | 0.1066 | 0.9621 | 0.1335 | 0.5476 | 0.0006 | 0.141 |
| Day 7 | Control vs. Fracture | 0.8914 | 0.6956 | 0.8934 | 0.7584 | 0.955 | 0.2245 | 0.9321 | 0.3598 | 0.4789 | 0.6675 | 0.8824 | 0.7357 | 0.2435 |
| | Control vs. MHV | 0.0002 | 0.0039 | 0.0271 | 0.0004 | 0.0002 | 0.04 | 0.0349 | 0.0457 | 0.0001 | 0.0089 | 0.864 | 0.2311 | 0.3026 |
| | Control vs. MHV + Fracture | 0.0899 | 0.1284 | 0.6709 | 0.7584 | 0.955 | 0.013 | 0.0984 | 0.0885 | 0.0002 | 0.3086 | 0.864 | 0.7198 | 0.5535 |
| | Fracture vs. MHV | 0.0003 | 0.0017 | 0.1236 | 0.0006 | 0.0007 | 0.0955 | 0.0349 | 0.0434 | 0.0003 | 0.16 | 0.864 | 0.2311 | 0.5535 |
| | Fracture vs. MHV + Fracture | 0.0899 | 0.1276 | 0.6709 | 0.7584 | 0.955 | 0.016 | 0.0984 | 0.0885 | 0.0002 | 0.2462 | 0.864 | 0.7198 | 0.6235 |
| | MHV vs. MHV + Fracture | 0.6719 | 0.6956 | 0.8934 | 0.7584 | 0.8444 | 0.2245 | 0.4804 | 0.3156 | 0.3411 | 0.6998 | 0.864 | 0.7198 | 0.7166 |

**Supplementary Table 4:** Male Flow Cytometry Holm-Sidak Pairwise Comparison (Time Point within Condition) (p-value)

| | | Lymphoid | B cells | T cells | CD8+ T cells | CD4+ T cells | CD25+ T cells | Myeloid | Monocytes | M1/M2 Monocytes | Macrophages | M2 Macrophages | Dendritic cells | Neutrophils |
|---|---|---|---|---|---|---|---|---|---|---|---|---|---|---|
| Control | Baseline vs Day 2 | 0.559 | 0.3905 | 0.5838 | 0.0062 | 0.0107 | 0.0148 | 0.1851 | 0.0003 | 0.0333 | 0.8101 | 0.0888 | 0.1745 | 0.2857 |
| | Baseline vs Day 7 | 0.2546 | 0.1997 | 0.1363 | 0.0599 | 0.1352 | 0.0089 | 0.5488 | 0.0003 | 0.0196 | 0.8101 | 0.0888 | 0.1745 | 0.3255 |
| | Day 2 vs Day 7 | 0.2546 | 0.2709 | 0.0899 | 0.1072 | 0.1352 | 0.3207 | 0.1851 | 0.7748 | 0.4526 | 0.8101 | 0.5756 | 0.0401 | 0.1971 |
| Fracture | Baseline vs Day 2 | 0.1995 | 0.9661 | 0.5742 | 0.0008 | 0.0036 | 0.0245 | 0.7152 | 0.0024 | 0.0165 | 0.4587 | 0.0931 | 0.3015 | 0.3834 |
| | Baseline vs Day 7 | 0.1572 | 0.9661 | 0.5742 | 0.1089 | 0.2175 | 0.019 | 0.7152 | 0.0003 | 0.0165 | 0.9391 | 0.0931 | 0.3377 | 0.9618 |
| | Day 2 vs Day 7 | 0.1122 | 0.9661 | 0.0164 | 0.0065 | 0.0105 | 0.4033 | 0.1298 | 0.0221 | 0.0881 | 0.0137 | 0.3145 | 0.3377 | 0.0296 |
| MHV | Baseline vs Day 2 | 0.0113 | 0.0133 | 0.0052 | 0.0929 | 0.8906 | 0.9368 | 0.0458 | 0.0004 | 0.0677 | 0.0495 | 0.1126 | 0.0034 | 0.246 |
| | Baseline vs Day 7 | 0.0113 | 0.0078 | 0.0022 | 0.0031 | 0.004 | 0.9489 | 0.0299 | 0.5219 | 0.0209 | 0.0594 | 0.1126 | 0.6412 | 0.5173 |
| | Day 2 vs Day 7 | 0.037 | 0.0186 | 0.0145 | 0.0031 | 0.0199 | 0.9489 | 0.1776 | 0.0177 | 0.0209 | 0.0026 | 0.8046 | <0.0001 | 0.7234 |
| MHV + Fracture | Baseline vs Day 2 | 0.0009 | 0.0207 | 0.006 | 0.0908 | 0.3643 | 0.219 | 0.2744 | 0.0003 | 0.3991 | 0.5455 | 0.143 | 0.6673 | 0.0054 |
| | Baseline vs Day 7 | 0.0539 | 0.1204 | 0.5321 | 0.5062 | 0.9617 | 0.219 | 0.1043 | 0.2847 | 0.1797 | 0.5455 | 0.4556 | 0.6673 | 0.4338 |
| | Day 2 vs Day 7 | 0.0277 | 0.6247 | 0.5321 | 0.5923 | 0.9617 | 0.6462 | 0.2451 | 0.2847 | 0.3991 | 0.1031 | 0.4556 | 0.6673 | 0.0241 |

**Supplementary Table 5:** Female Multiplex Holm-Sidak Pairwise Comparison (Condition within Time Point) (p-value)

| | | IL-18 | GRO | TNFα | IP-10 | BAFF | IL-2R | VEGF-A | Leptin | CXCL2 |
|---|---|---|---|---|---|---|---|---|---|---|
| Baseline | Control vs. Fracture | >0.9999 | >0.9999 | >0.9999 | >0.9999 | >0.9999 | >0.9999 | >0.9999 | >0.9999 | >0.9999 |
| | Control vs. MHV | >0.9999 | >0.9999 | >0.9999 | >0.9999 | >0.9999 | >0.9999 | >0.9999 | >0.9999 | >0.9999 |
| | Control vs. MHV + Fracture | >0.9999 | >0.9999 | >0.9999 | >0.9999 | >0.9999 | >0.9999 | >0.9999 | >0.9999 | >0.9999 |
| | Fracture vs. MHV | >0.9999 | >0.9999 | >0.9999 | >0.9999 | >0.9999 | >0.9999 | >0.9999 | >0.9999 | >0.9999 |
| | Fracture vs. MHV + Fracture | >0.9999 | >0.9999 | >0.9999 | >0.9999 | >0.9999 | >0.9999 | >0.9999 | >0.9999 | >0.9999 |
| | MHV vs. MHV + Fracture | >0.9999 | >0.9999 | >0.9999 | >0.9999 | >0.9999 | >0.9999 | >0.9999 | >0.9999 | >0.9999 |
| Day 2 | Control vs. Fracture | 0.5369 | 0.5645 | 0.5339 | 0.3872 | 0.7553 | 0.188 | 0.6433 | 0.7613 | 0.6591 |
| | Control vs. MHV | 0.3753 | 0.2736 | 0.1562 | 0.1911 | 0.7553 | 0.188 | 0.0465* | 0.8332 | 0.6591 |
| | Control vs. MHV + Fracture | 0.0042* | 0.2051 | 0.0158* | 0.0265* | 0.1789 | 0.0759 | 0.0465* | 0.4298 | 0.6591 |
| | Fracture vs. MHV | 0.1585 | 0.7561 | 0.1609 | 0.0861 | 0.8836 | 0.6709 | 0.1217 | 0.8332 | 0.6591 |
| | Fracture vs. MHV + Fracture | 0.0029* | 0.5645 | 0.0158* | 0.0233* | 0.1362 | 0.0759 | 0.1217 | 0.6573 | 0.6591 |
| | MHV vs. MHV + Fracture | 0.0125* | 0.5645 | 0.1609 | 0.1911 | 0.1789 | 0.188 | 0.6433 | 0.6573 | 0.6591 |
| Day 7 | Control vs. Fracture | 0.9995 | 0.5152 | 0.8854 | 0.9886 | 0.4819 | 0.2 | 0.8387 | 0.6771 | 0.8377 |
| | Control vs. MHV | 0.9995 | 0.9353 | 0.8854 | 0.2021 | 0.2025 | 0.0951 | 0.1929 | 0.191 | 0.477 |
| | Control vs. MHV + Fracture | 0.9922 | 0.9353 | 0.8854 | 0.9886 | 0.0144* | 0.0632 | 0.2564 | 0.191 | 0.477 |
| | Fracture vs. MHV | 0.9995 | 0.5152 | 0.8854 | 0.1359 | 0.1167 | 0.2 | 0.0205* | 0.0225* | 0.3382 |
| | Fracture vs. MHV + Fracture | 0.9922 | 0.9043 | 0.8854 | 0.9886 | 0.0175* | 0.0951 | 0.2554 | 0.0081* | 0.3382 |
| | MHV vs. MHV + Fracture | 0.9922 | 0.9353 | 0.8854 | 0.7789 | 0.4819 | 0.4644 | 0.8387 | 0.6844 | 0.8377 |

**Supplementary Table 6:** Female Multiplex Holm-Sidak Pairwise Comparison (Time Point within Condition) (p-value)

| | | IL-18 | GRO | TNFα | IP-10 | BAFF | IL-2R | VEGF-A | Leptin | CXCL2 |
|---|---|---|---|---|---|---|---|---|---|---|
| Control | Baseline vs Day 2 | 0.8244 | 0.5393 | 0.4725 | 0.7563 | 0.9982 | 0.9639 | 0.7123 | 0.9589 | 0.8531 |
| | Baseline vs Day 7 | 0.9442 | 0.6636 | 0.9307 | 0.2899 | 0.9982 | 0.9393 | 0.7123 | 0.9589 | 0.963 |
| | Day 2 vs Day 7 | 0.7139 | 0.5393 | 0.4782 | 0.2899 | 0.9982 | 0.7372 | 0.7123 | 0.9104 | 0.8531 |
| Fracture | Baseline vs Day 2 | 0.6478 | 0.4944 | 0.3713 | 0.6765 | 0.6756 | 0.4657 | 0.7701 | 0.8158 | 0.8066 |
| | Baseline vs Day 7 | 0.9729 | 0.2204 | 0.8631 | 0.1032 | 0.6304 | 0.3797 | 0.6965 | 0.8158 | 0.8511 |
| | Day 2 vs Day 7 | 0.4801 | 0.457 | 0.8965 | 0.1032 | 0.4622 | 0.9644 | 0.7701 | 0.6385 | 0.6798 |
| MHV | Baseline vs Day 2 | N/A | 0.3092 | 0.6643 | 0.2326 | 0.6953 | 0.2522 | 0.5064 | 0.5513 | 0.7475 |
| | Baseline vs Day 7 | N/A | 0.8308 | 0.7764 | 0.4752 | 0.4455 | 0.2522 | 0.5064 | 0.3385 | 0.7475 |
| | Day 2 vs Day 7 | N/A | 0.127 | 0.9161 | 0.2326 | 0.3646 | 0.0182* | 0.5064 | 0.3385 | 0.7475 |
| MHV + Fracture | Baseline vs Day 2 | N/A | 0.1498 | 0.2879 | 0.0715 | 0.2577 | 0.266 | 0.5017 | 0.5678 | 0.7582 |
| | Baseline vs Day 7 | N/A | 0.4638 | 0.8311 | 0.5585 | 0.1905 | 0.266 | 0.6315 | 0.3619 | 0.7582 |
| | Day 2 vs Day 7 | N/A | 0.1655 | 0.8364 | 0.6432 | 0.0734 | 0.8319 | 0.6315 | 0.638 | 0.7582 |

**Supplementary Table 7:** Male Multiplex Holm-Sidak Pairwise Comparison (Condition within Time Point) (p-value)

| | | IL-2R | IL-2 | Leptin | BAFF | CCL7 | IL-10 | IL-4 | VEGF-A |
|---|---|---|---|---|---|---|---|---|---|
| Baseline | Control vs. Fracture | >0.9999 | >0.9999 | >0.9999 | >0.9999 | >0.9999 | >0.9999 | >0.9999 | >0.9999 |
| | Control vs. MHV | >0.9999 | >0.9999 | >0.9999 | >0.9999 | >0.9999 | >0.9999 | >0.9999 | >0.9999 |
| | Control vs. MHV + Fracture | >0.9999 | >0.9999 | >0.9999 | >0.9999 | >0.9999 | >0.9999 | >0.9999 | >0.9999 |
| | Fracture vs. MHV | >0.9999 | >0.9999 | >0.9999 | >0.9999 | >0.9999 | >0.9999 | >0.9999 | >0.9999 |
| | Fracture vs. MHV + Fracture | >0.9999 | >0.9999 | >0.9999 | >0.9999 | >0.9999 | >0.9999 | >0.9999 | >0.9999 |
| | MHV vs. MHV + Fracture | >0.9999 | >0.9999 | >0.9999 | >0.9999 | >0.9999 | >0.9999 | >0.9999 | >0.9999 |
| Day 2 | Control vs. Fracture | 0.1443 | 0.0884 | 0.0072 | 0.0442 | 0.6442 | N/A | 0.6177 | 0.9602 |
| | Control vs. MHV | 0.0627 | 0.1133 | 0.0072 | 0.6591 | 0.5873 | N/A | 0.9653 | 0.1121 |
| | Control vs. MHV + Fracture | 0.0433 | 0.1133 | 0.0072 | 0.0155 | 0.0925 | N/A | 0.9198 | 0.9602 |
| | Fracture vs. MHV | 0.3123 | 0.4268 | 0.2236 | 0.0442 | 0.5873 | N/A | 0.0876 | 0.8002 |
| | Fracture vs. MHV + Fracture | 0.2092 | 0.4286 | 0.2236 | 0.6591 | 0.0332 | N/A | 0.0697 | 0.9673 |
| | MHV vs. MHV + Fracture | 0.1044 | 0.7404 | 0.9658 | 0.0116 | 0.2192 | N/A | 0.7999 | 0.9602 |
| Day 7 | Control vs. Fracture | 0.4072 | N/A | 0.9044 | 0.6866 | 0.9782 | 0.4228 | 0.562 | 0.4183 |
| | Control vs. MHV | 0.0263 | N/A | 0.9044 | 0.0149 | 0.1791 | 0.3343 | 0.3342 | 0.2719 |
| | Control vs. MHV + Fracture | 0.0263 | N/A | 0.5334 | 0.0149 | 0.493 | 0.3343 | 0.562 | 0.1192 |
| | Fracture vs. MHV | 0.2677 | N/A | 0.9044 | 0.0144 | 0.1852 | 0.4228 | 0.0005 | 0.3361 |
| | Fracture vs. MHV + Fracture | 0.266 | N/A | 0.9044 | 0.0149 | 0.493 | 0.4228 | 0.1849 | 0.2901 |
| | MHV vs. MHV + Fracture | 0.4072 | N/A | 0.9982 | 0.6866 | 0.579 | 0.9725 | 0.562 | 0.3361 |

**Supplementary Table 8:** Male Multiplex Holm-Sidak Pairwise Comparison (Time Point within Condition) (p-value)

| | | IL-2R | IL-2 | Leptin | BAFF | CCL7 | IL-10 | IL-4 | VEGF-A |
|---|---|---|---|---|---|---|---|---|---|
| Control | Baseline vs Day 2 | 0.6666 | 0.7889 | 0.6035 | 0.8989 | 0.895 | 0.8039 | 0.6185 | 0.2248 |
| | Baseline vs Day 7 | 0.7649 | 0.9932 | 0.6035 | 0.8989 | 0.8377 | 0.1515 | 0.8445 | 0.4151 |
| | Day 2 vs Day 7 | 0.209 | 0.7889 | 0.6042 | 0.8989 | 0.895 | 0.8902 | 0.6196 | 0.4151 |
| Fracture | Baseline vs Day 2 | 0.6719 | 0.229 | 0.2991 | 0.4174 | 0.785 | N/A | 0.745 | 0.628 |
| | Baseline vs Day 7 | 0.6719 | 0.4931 | 0.3103 | 0.9853 | 0.785 | N/A | 0.3484 | 0.4876 |
| | Day 2 vs Day 7 | 0.9968 | 0.4931 | 0.2991 | 0.081 | 0.8145 | N/A | 0.9272 | 0.7843 |
| MHV | Baseline vs Day 2 | 0.1771 | 0.6557 | 0.2472 | 0.8653 | 0.653 | 0.3102 | 0.0071 | 0.7264 |
| | Baseline vs Day 7 | 0.1771 | 0.6557 | 0.7823 | 0.1655 | 0.653 | 0.3102 | 0.0624 | 0.7264 |
| | Day 2 vs Day 7 | 0.0645 | 0.2641 | 0.58 | 0.0007 | 0.0178 | 0.7408 | 0.3147 | 0.7264 |
| MHV + Fracture | Baseline vs Day 2 | 0.2233 | N/A | 0.4321 | 0.2719 | 0.1839 | 0.3297 | 0.2736 | 0.6888 |
| | Baseline vs Day 7 | 0.2233 | N/A | 0.4321 | 0.1291 | 0.698 | 0.3297 | 0.3482 | 0.2742 |
| | Day 2 vs Day 7 | 0.411 | N/A | 0.4321 | 0.0279 | 0.0586 | 0.9291 | 0.5869 | 0.6888 |
